# Virus and CTL dynamics in the extra-follicular and follicular tissue compartments in SIV-infected macaques

**DOI:** 10.1101/330217

**Authors:** Dominik Wodarz, Pamela J. Skinner, David N. Levy, Elizabeth Connick

## Abstract

Data from SIV-infected macaques indicate that virus-specific cytotoxic T lymphocytes (CTL) are mostly present in the extrafollicular (EF) compartment of the lymphoid tissue, with reduced homing to the follicular (F) site. This contributes to the majority of the virus being present in the follicle and represents a barrier to virus control. Using mathematical models, we investigate these dynamics. Two models are analyzed. The first assumes that CTL can only become stimulated and expand in the extra-follicular compartment, with migration accounting for the presence of CTL in the follicle. In the second model, follicular CTL can also undergo antigen-induced expansion. Consistent with experimental data, both models predict increased virus compartmentalization in the presence of stronger CTL responses and lower virus loads, and a more pronounced rise of extra-follicular compared to follicular virus during CD8 cell depletion experiments. The models, however, differ in other aspects. The follicular expansion model results in dynamics that promote the clearance of productive infection in the extrafollicular site, with any productively infected cells found being the result of immigration from the follicle. This is not observed in the model without follicular CTL expansion. The models further predict different consequences of introducing engineered, follicular-homing CTL, which has been proposed as a therapeutic means to improve virus control. Without follicular CTL expansion, this is predicted to result in a reduction of virus load in both compartments. The follicular CTL expansion model, however, makes the counter-intuitive prediction that addition of F-homing CTL not only results in a reduction of follicular virus load, but also in an increase in extrafollicular virus replication. These predictions remain to be experimentally tested, which will be relevant for distinguishing between models and for understanding how therapeutic introduction of F-homing CTL might impact the overall dynamics of the infection.

## Introduction

Human immunodeficiency virus (HIV-1) causes a persistent infection that eventually progresses to AIDS, following an asymptomatic phase that is variable in duration. Immune responses have been shown to play a major role in determining the level of virus load and the rate of disease progression [1–4]. Cytotoxic T lymphocytes (CTL) are an important branch of the immune system in this respect [1,5,6]. The dynamics between the virus and CTL responses have been documented largely in the blood, and a variety of insights have been obtained about the role of CTL responses for virus control in HIV-infected patients [1,7]. The SIV-infected rhesus macaque model recapitulates many aspects of HIV immunopathogenesis including the induction of virus-specific CTL responses.

The majority of HIV-1 and SIV replication occurs in secondary lymphoid tissues including lymph nodes, spleen, and mucosal associated lymphoid tissues [8–10]. Within secondary lymphoid tissues, the dynamics of virus replication are influenced by the CTL response. CTL are generated in the extra-follicular compartment and display limited homing to the follicular compartment in both HIV-infected humans [11] and SIV-infected rhesus macaques [12]. In SIV-infected macaques with robust CTL responses, virus replication is generally low and the majority of replication is located in the follicular compartment [12]. In contrast, animals with ineffective or absent CTL responses display much more virus replication and an equal distribution of infected cells in the two compartments [12]. These observations give rise to the notion that the presence of effective CTL responses (leading to relatively low virus loads) contributes to the observed unequal distribution of the virus in the two compartments [13].

Here, we investigate these dynamics with the help of two-compartment mathematical models. In agreement with published and newly presented data, the models show that variation in the strength of the CTL response can influence the degree of virus compartmentalization, with stronger CTL responses (that result in lower virus load) correlating with a more unequal distribution of the virus population among the two compartments. Interestingly, details of the outcome of these dynamics can depend on assumptions about CTL activity in the follicular compartment. If it is assumed that CTL can be stimulated and expand to an extent in the follicular compartment, the model predicts a relatively low number of infected cells in the extra-follicular compartment, maintained mostly by immigration from the follicle rather than by successful replication in the extra-follicular compartment itself. If, in contrast, the model assumes that follicular CTL cannot be stimulated to expand, an unequal virus distribution among the compartments still occurs in the model, but higher virus loads are predicted for the extra-follicular compartment, maintained by successful and sustained viral replication in this site rather than by immigration from the follicle. The models further suggest that future experiments involving the addition of follicular-homing CTL to SIV-infected macaques could distinguish between these two assumptions. Without follicular CTL expansion, addition of F-homing CTL is predicted to result in a significant reduction of follicular virus load, and a lesser reduction of extrafollicualr virus load. In contrast, if CTL expansion is allowed to occur, the reduction of follicular virus load upon addition of F-homing CTL is predicted to be accompanied by a significant rise of extrafollicular virus load. A better understanding of these dynamics also has implications for potentially developing the therapeutic use of chimeric antigen receptor T cells that express the follicular homing molecule CXCR5 to improve the overall degree of virus control.

## Results

### A basic two-compartment model without CTL stimulation in the follicular compartment

We consider a two-compartment mathematical model for virus replication (Figure 1A). The virus population can replicate either in the follicular compartment, or in the extra-follicular compartment. The number of uninfected and infected cells in the follicular compartment are denoted by X_f_ and Y_f_, respectively. The corresponding populations in the extra-follicular compartment are denoted by X_e_ and Y_e_, respectively. The main CTL dynamics are assumed to occur in the extralfollicular compartment, and this population is denoted by Z_e_. CTL are also assumed to be able to enter the follicular compartment, and the CTL population there is denoted by Z_f_. The model is thus given by the following set of ordinary differential equations, which describe the average time evolution of the populations.

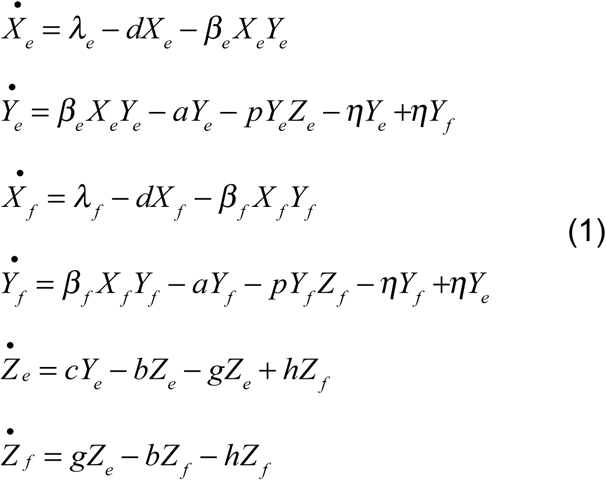

**Figure 1.**
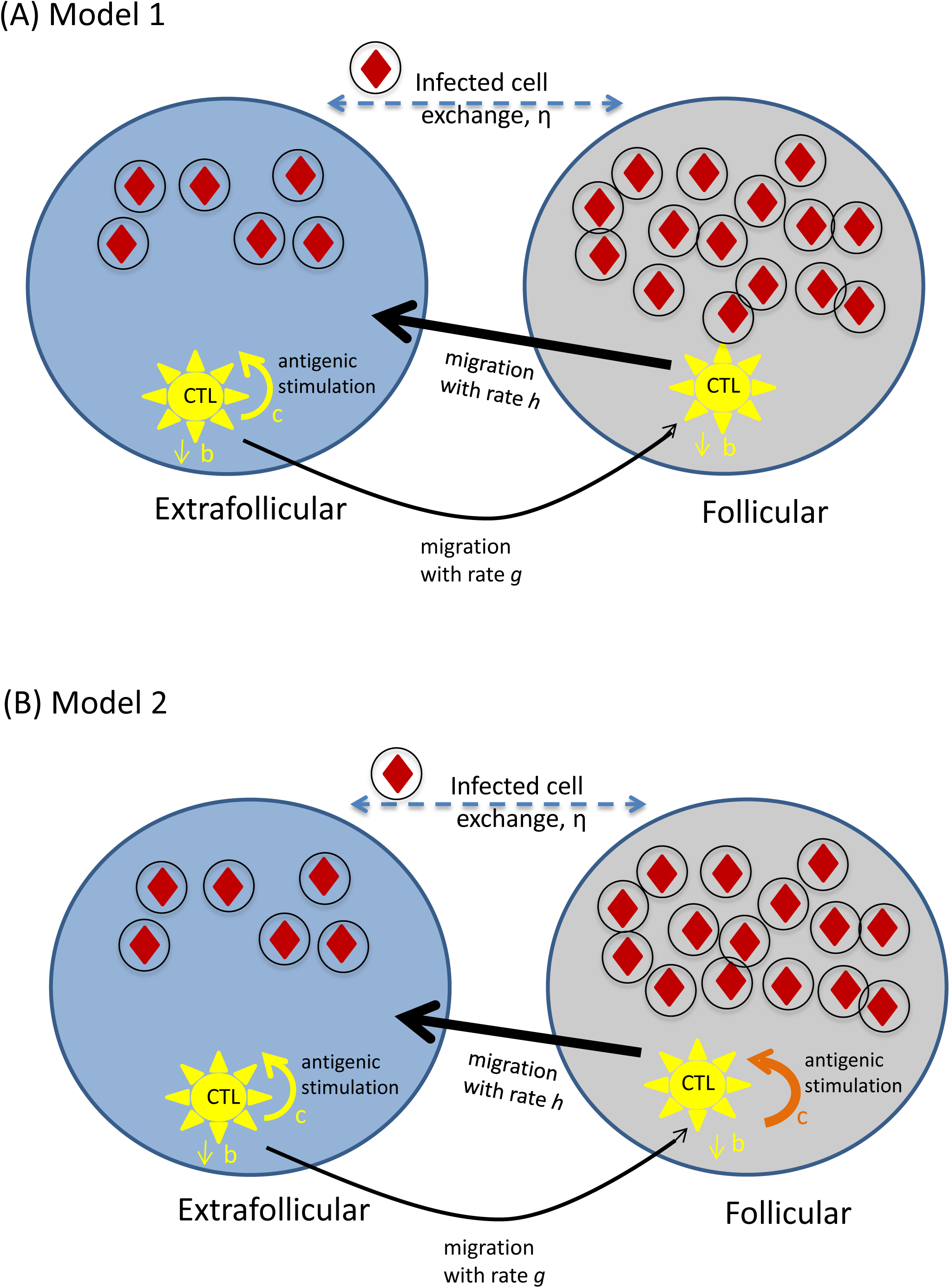
Schematic representation of key model assumptions. (A) Model (1) assuming absence of antigen-induced CTL stimulation and expansion in the follicular compartment. (B) Model (2) assuming that CTL can become stimulated and expand in the follicular compartment. Infected cells are depicted by circles including red diamonds. CTL are shown in yellow. The difference between the two models solely lies in the ability of CTL to expand in the follicular compartment, which is indicated by the absence / presence of the orange expansion arrow in parts A/B.

These equations are an extension of the well-established theoretical framework to mathematically describe the dynamics of virus infections [14–19]. The model does not explicitly take into account the free virus population, which is assumed to be in a quasisteady state with infected cells (i.e. proportional to the number of infected cells [14]). First, consider the extrafollicular compartment. Uninfected cells are produced with a rate λ_e_, die with a rate d, and upon contact with virus are infected with a rate β_e_. Infected cells die with a rate a, and are killed by CTL with a rate p_e_. The same basic virus dynamics occur in the follicular compartment. Thus, uninfected cells are produced with a rate λ_f_, die with a rate d, and upon contact with virus are infected with a rate β_f_. Infected cells die with a rate a, and are killed by CTL with a rate p_f_. We assume that infected cells can migrate between the two compartments with a rate η. The dynamics of CTL expansion only occur in the extrafollicular compartment. Hence, the CTL expand in response to antigenic stimulation with a rate c. In the absence of stimulation, CTL are assumed to die with a rate b. Extrafolliclar CTL can migrate to the follicular compartment with a rate g. In the follicular compartment, no CTL stimulation / expansion is assumed to occur. CTL can die with a rate b, and they can migrate back to the extrafollicular compartment with a rate h. We note that this model does not assume migration of uninfected cells between the two compartments. A low migration rate would not change the model properties in a meaningful way. If uninfected cells moved between the two compartments with a fast rate, this would affect the relative abundance of the uninfected cells in the two compartments. As mentioned below in Section 5, the distribution of target cells among the two compartments has not been measured, and there is no significant change in conclusions if it is varied within reason. We further note that instead of explicitly tracking the free virus population, we assumed it to be in a quasi-steady state, proportional to the number of infected cells [14]. While it might be possible for free viruses to diffuse more readily between the two compartments than infected cells, fast virus exchange would equalize the distribution of the viruses across the compartments even in the presence of strong immune responses, which is contrary to observations. Hence, the quasi-steady state assumption is likely sufficient for our analysis.

Whether an infection can be established in the absence of immune responses is given by the basic reproductive ratio of the virus [14]. This is defined as the average number of newly infected cells produced by one infected cell during its lifespan when placed into a pool of susceptible cells. It is instructive to consider the situation where there is no virus migration between the two compartments (η=0), because this significantly simplifies the expressions. In this case, we can define the basic reproductive ratio separately for the follicular and the extra-follicular compartment, given by R_0f_ = λ_f_β_f_/da and R_0e_ = λ_e_β_e_/da. If R_0f_>1, the virus establishes an infection in the follicular compartment. If R_0e_>1, the virus establishes an infection in the extrafollicular compartment. If these conditions are fulfilled, the populations converge to the following equilibrium in the absence of immunity.

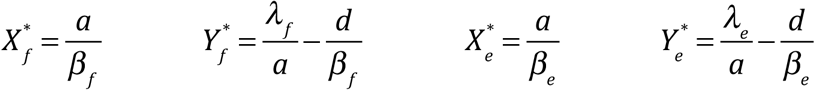

Returning to the biologically realistic scenario where infected cell migration occurs between the two compartments (η>0), the virus establishes an infection if the following condition is fulfilled: (*ad* − *β* _*f*_*λ* _*f*_)(*ad* − *β*_*e*_*λ*_*e*_)+ *dη*(2*ad* − *β* _*f*_*λ* _*f*_− *β*_*e*_*λ*_*e*_) > 0. In this case, the system converges to an equilibrium, the expressions for which are too lengthy to write down here. For low values of η, however, the expressions converge to the ones derived for η=0 above. When a CTL response is added to the system, it reduces virus load towards an equilibrium that is again too lengthy and uninformative to write down. In the mathematical formulation used here, the CTL response always becomes established if the infection persists in the absence of immunity [17] (which is in contrast to other model formulations where establishment of CTL requires a threshold virus load [17]).

In computer simulations, parameter values need to be chosen. Several parameters are unknown (especially for immune responses and compartment-specific processes), and they were chosen arbitrarily for the purpose of illustration. The presented simulation results are not dependent on this particular choice of parameter values unless otherwise stated (see section 5 for more in depth discussion of parameter effects). Known parameters are adopted from the literature. Hence, infected cells are assumed to have an average life-span of around 2.2 days [20], and other viral replication parameters were adjusted such that the basic reproductive ratio of the virus was approximately eight [21–23]. In the following, we examine how immune parameters determine virus load in the two compartments. We will thereby concentrate on two parameters. The first is the strength of the CTL response, defined as the rate at which CTL respond to antigen, or CTL responsiveness c. This parameter captures the many biological processes that contribute to the rate of activation and expansion when a CTL is exposed to antigenic stimulation. The second parameter is the rate of CTL migration from the extrafollicular to the follicular compartment, g. While we concentrate on those parameters, the effect of varying other parameters, as well as scaling effects, are described in Section 5.

#### Variation in the CTL responsiveness, c

In the mathematical model considered, the CTL responsiveness parameter, c, is one of the most important immune parameters that determines virus load at equilibrium [24]. Thus, a lower value of c results in higher virus load, which can correspond to advanced stages of the disease when immune responses have been weakened. Similar considerations could apply to the very early stages of the disease, before immune responses have fully developed, although this does not represent any kind of equilibrium situation. A higher value of c would correspond to the early chronic phase of the infection, especially in good controllers, when virus load is relatively low. In the following, we assume that virus infection parameters are the same in both compartments. Figure 2A plots the equilibrium number of infected cells in the two compartments, as a function of the CTL responsiveness, c. An increase in the parameter c results in overall lower virus loads. The decline in virus load is more pronounced in the extrafollicular compared to the follicular compartment (Figure 2A). For very large values of c, the equilibrium number of extrafollicular infected cells tends towards zero, while the equilibrium number of follicular infected cells tends towards a constant. These trends is also reflected in Figure 2B, showing a negative association between total virus load and the ratio of follicular : extra-follicular (F:EF) virus load, as the rate of CTL expansion is varied. That is, the lower the overall number of infected cells, the stronger the degree of compartmentalization observed in the model. For the highest virus loads in the model (weak CTL), the distribution of the infected cell populations in the two compartments becomes equal for the chosen parameters (F:EF ratio converges to 1, Figure 2A&B). This is consistent with experimental data that show a dominance of the virus population in the follicular compartment in the chronic phase of SIV infection, when CTL responses are strong, and a more equal distribution during advanced disease or in the very early stages of infection, when CTL-mediated suppression of the virus is weaker [12].

**Figure 2.**
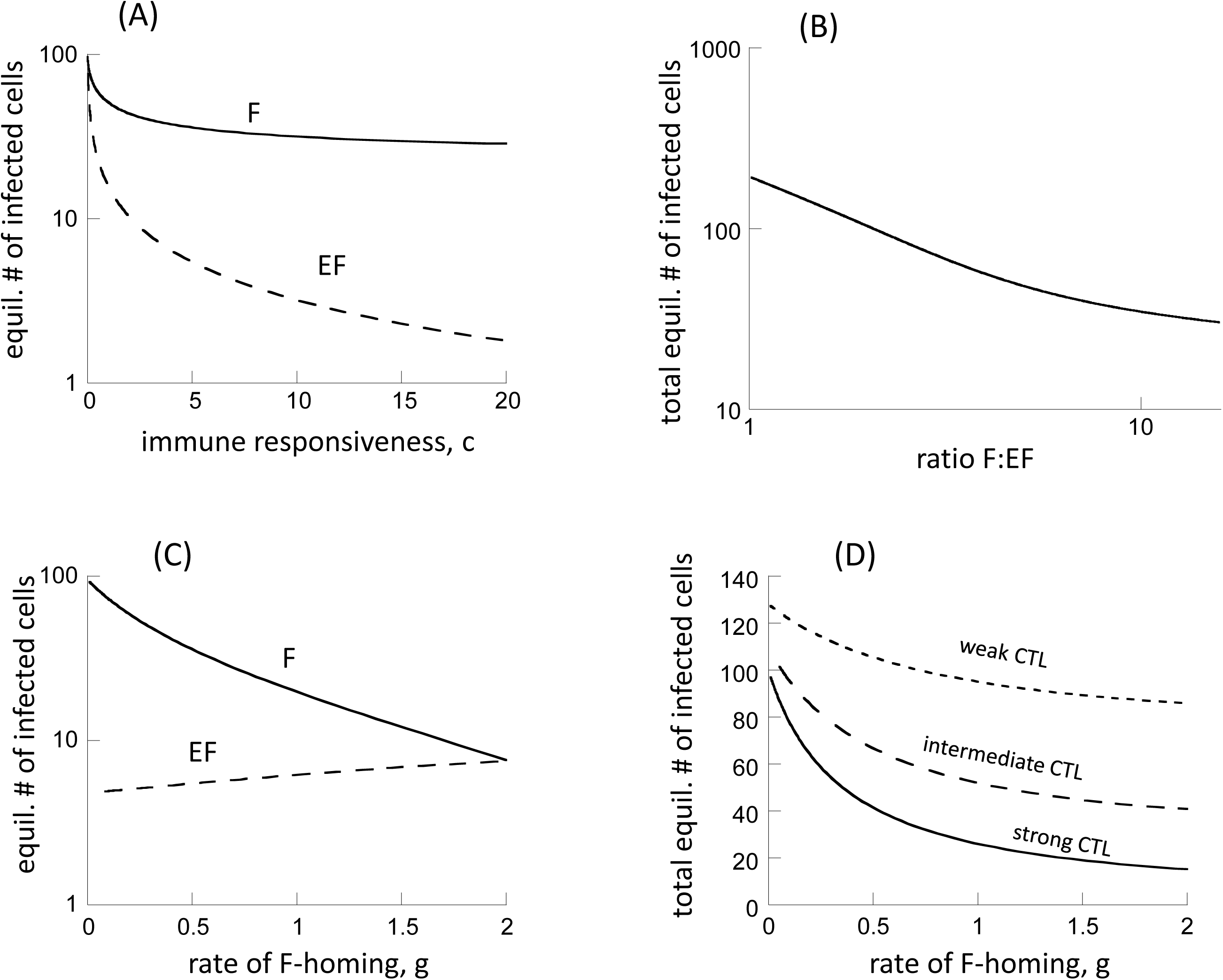
Properties of model (1), assuming absence of antigen-induced stimulation in the follicular compartment. (A) Equilibrium number of virus-producing cells in the two compartments as a function of the CTL responsiveness, c, which correlates with the strength of the CTL response. F=follicular compartment, EF=extrafollicular compartment. (B) Relationship between the equilibrium number of virus-producing cells and the ratio of follicular to extra-follicular virus load (F:EF). (C) Equilibrium number of virusproducing cells in the two compartments as a function of the rate at which CTL home to the follicular compartment, g. We note that the graph does not start at g=0, but at g=0.01. (D) Total equilibrium number of virus-producing cells (summed over both compartments) as a function of the rate at which CTL home to the follicular compartment, g. Base parameters were chosen as follows. λ_e_= λ_f_ =50, d=0.01, β_e_=, β_f_=0.00072, a=0.45, η=0.01, c_e_=5, b=0.01, p=0.001, g=0.5, h=2. For (D), we assumed c_e_=5 for the strong CTL response, c_e_=1 for the intermediate response, and c_e_=0.2 for the weak response.

#### Variation in the rate of CTL homing to the follicular compartment, g

The rate at which CTL move from the extrafollicular to the follicular compartment (g) can also influence the distribution of virus in the two compartments in the model. It is unclear whether this parameter is biologically fixed or whether it can be influenced by the virus. Varying this parameter in the model, however, allows us to study how it can potentially affect infection outcome. At relatively low CTL F-homing rates, the virus is more abundant in the F compared to the EF compartment (Figure 2C). If more CTL move from EF to F (larger value of g), virus load in the F compartment decreases significantly, because more CTL-mediated killing occurs in the F compartment (Figure 2C). At the same time, virus load in the EF compartment rises a little (Figure 2C), because homing of CTL to the follicular compartment leaves fewer CTL in the extrafollicular compartment. This rise, however, is not very pronounced. If the homing rate of CTL from the extra-follicular to the follicular compartment becomes equal to the movement of CTL back from the F to the EF compartment (g=2, Figure 2C), the distribution of the virus population across the two compartments becomes even for the parameter combinations considered. Summed over both compartments, total virus load declines with increased homing of CTL to the F compartment (Figure 2D). The extent of this decline, however, depends on the strength of the CTL response (parameter c). The stronger the CTL response, the more pronounced this effect (Figure 2D).

### Model with CTL stimulation in the follicular compartment

Here, the above analysis is repeated, assuming that CTL can also be stimulated and expand in the follicular compartment (Figure 1B). The equations for the CTL dynamics are thus given by

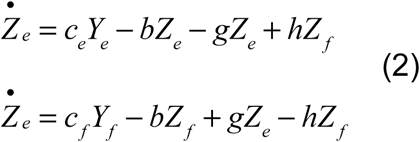

The equations for the virus dynamics remain the same as in model (1). The CTL are now assumed to expand in both compartments, with rates c_e_ and c_f_, respectively. Since we currently do not have any information about the relative magnitude of these parameters, we will assume c= c_e_ = c_f_ for purposes of illustration. Results, however, do not depend on this particular choice (see section 5). Although CTL in model (2) can expand in response to antigenic stimulation in the follicular compartment, the rate at which they migrate back to the extrafollicular compartment is set to be relatively large. The effect of this is that while stimulation does occur in the follicle, most of the cells that arise from this expansion quickly re-enter the extrafollicular compartment, thus limiting the number of CTL that remain in the follicle. In the absence of this assumption, follicular CTL expansion would result in accumulation of CTL at this location and would thus be unrealistic.

While this model generally has similar properties as model (1), there are some significant differences that arise from the assumption that CTL can be stimulated in the follicular compartment (and subsequently re-enter the EF compartment to exert immune pressure there). This introduces an element of “indirect” or “apparent” competition among viruses in the different locations [25,26], mediated by a shared CTL response. In model (1), the dynamics in the extrafollicular compartment were governed by the interactions between the local virus population and the CTL population that was stimulated in that compartment only. In model (2), however, CTL are independently added to the extrafollicular compartment by immigration, following antigenic stimulation in the follicular compartment. This increases immunological pressure on the extrafollicuclar virus population. As a consequence, equilibrium virus load in the extrafollicular compartment is predicted to decline to a larger extent if the strength of the CTL response is increased, compared to model (1), see Figure 3A (we varied the rate of CTL expansion in both compartments, such that c=c_e_=c_f_). Similar to model (1), an increase in the F:EF ratio is observed as virus load becomes lower (Figure 3B). For the same parameter ranges as in model (1), however, we observe higher F:EF ratios in model (2) (Figure 3B). The reason is the more pronounced reduction in the equilibrium number of extrafollicular infected cells with stronger CTL responses, which leads to more extensive compartmentalization. The exact values of the F:EF ratios depends on the choices of parameter values, which are currently unknown.

**Figure 3.**
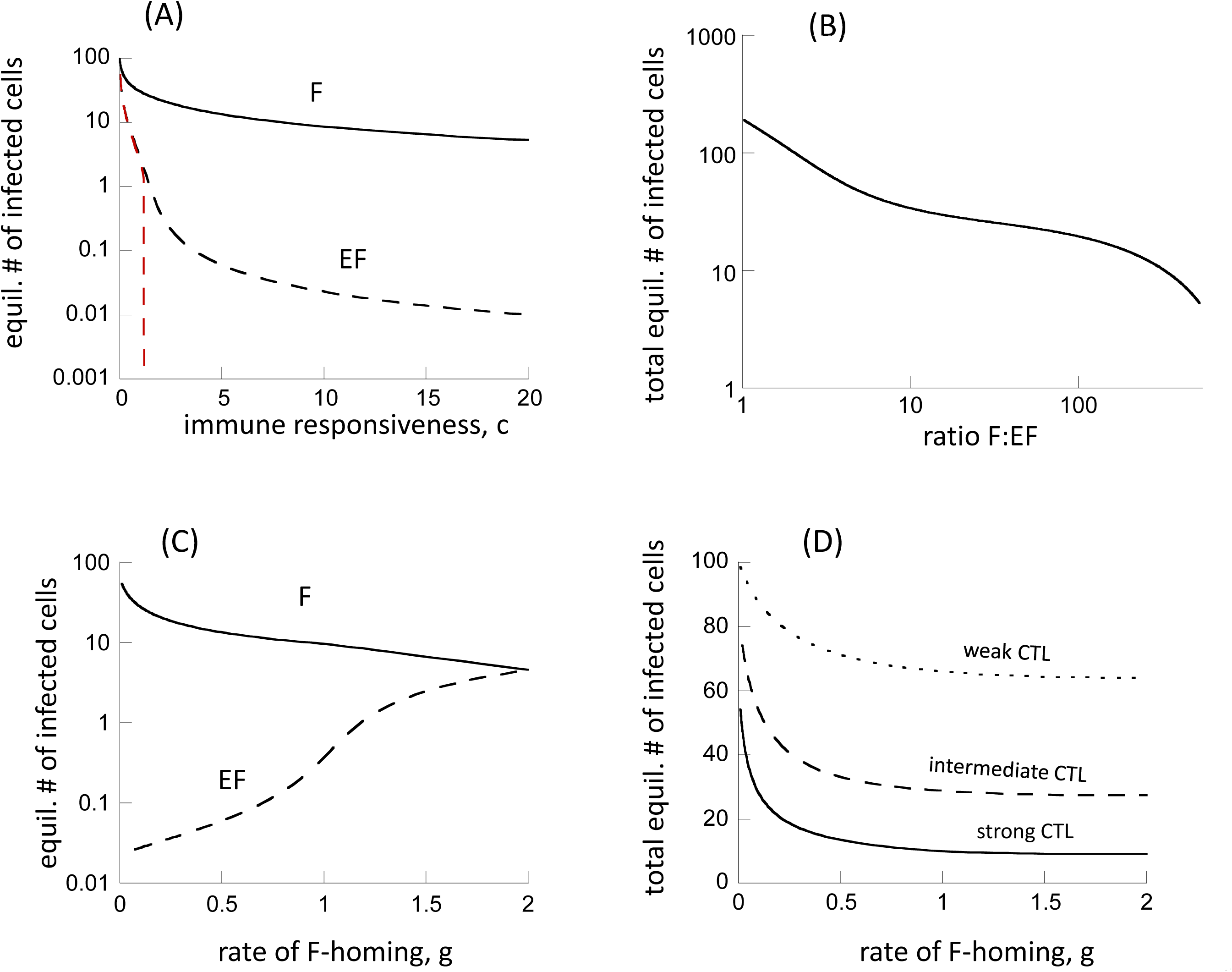
Properties of model (2), assuming the presence of antigen-induced stimulation in the follicular compartment. (A) Equilibrium number of virus-producing cells in the two compartments as a function of the rate of CTL expansion. Since CTL are assumed to expand in both compartments in this model, we varied the expansion rate in both, by setting c=c_e_=c_f_. The red dashed lined depicts results under the assumption that virus-producing cells cannot migrate between the two comparmtents, η=0. (B) Relationship between the equilibrium number of virus-producing cells and the ratio of follicular to extra-follicular virus load (F:EF). (C) Equilibrium number of virus-producing cells in the two compartments as a function of the rate at which CTL home to the follicular compartment, g. We note that the graph does not start at g=0, but at g=0.01. (D) Total equilibrium number of virus-producing cells (summed over both compartments) as a function of the rate at which CTL home to the follicular compartment, g. Base parameters were chosen as follows. λ_e_= λ_f_ =50, d=0.01, β_e_=, β_f_=0.00072, a=0.45, η=0.01, c_e_=c_f_=5, b=0.01, p=0.001, g=0.5, h=2. For (D), we assumed c_e_=5 for the strong CTL response, c_e_=1 for the intermediate response, and c_e_=0.2 for the weak response.

It is instructive to also plot the extrafollicular number of infected cells at equilibrium under the assumption that infected cells do not migrate between the two compartments (η=0, dashed red line in Figure 3A). Now, the extrafollicular number of virus-producing cells goes extinct if the strength of the CTL response, c, crosses a threshold. The reason is that the CTL that are stimulated in the F compartment and move to the EF compartment put extra pressure on the virus population in the EF compartment, thus clearing the virus in this site. In the presence of infected cell migration (η>0, dashed black line in Figure 3A), the persistence of extrafollicular virus beyond this CTL strength threshold is the result of infected cell migration from the follicular to the extrafollicular compartment, and not maintained by active virus replication in the EF compartment itself.

Some differences are also observed when varying the rate at which CTL home from the extrafollicular to the follicular compartment (parameter g, Figure 3C). If the rate of CTL homing to F is low (low g), there is relatively high virus load in the F compartment, and very few infected cells are present in the EF compartment, only due to immigration of infected cells from the follicular compartment, as described above. For higher rates of CTL homing to the F compartment, follicular virus is reduced because more CTL enter this compartment and kill infected cells. At the same time, virus load in the EF compartment rises significantly (much more than in model (1)), and is now maintained by ongoing productive infection in the EF compartment (rather than just by immigration). The reason is two-fold. (i) As in model (1), a larger number of the CTL leave the EF compartment and enter the F compartment, resulting in less killing of infected cells in the EF compartment. (ii) In addition, and more importantly, lower follicular virus load results in less stimulation of follicular CTL that can migrate back to the EF site, thus reducing external CTL-mediated pressure on EF virus. For large values of g that are comparable to the rate of CTL migration back to the EF compartment, h, the virus in the two compartments becomes equally distributed. While increased CTL homing to the follicular compartment is predicted to result in an increase in EF virus load and a decline in follicular virus load, total virus load (the sum of the virus population in F and EF compartments) is predicted to decline to a certain extent (Figure 3D). As in model (1), this decline becomes less pronounced for weaker CTL responses (Figure 3D).

### Predictions about experimental manipulations

In the above sections, we have shown how the dynamics of CTL responses can influence the distribution of virus load across the two compartments. Hence, experimental manipulation of the CTL responses, such as CTL depletion or addition, should result in changes in these distributions. We have considered two mathematical models, one assuming that CTL cannot be stimulated to proliferate in the follicular compartment, while the second model did allow for CTL stimulation in this site. Here, we explore how CTL depletion and addition experiments are predicted by the two models to change the virus distribution in the two compartments, and how these predictions compare to experimental data. We will explore whether the comparison between models and data can be used to distinguish between the two models considered.

#### Simulating CTL depletion experiments

Here we show model predictions about the outcome of experiments in which CTL are depleted from SIV-infected macaques, and compare predictions to experimental data. To do so, we compare the equilibrium number of infected cells in the presence of CTL (corresponding to pre-depletion), and in the absence of CTL (corresponding to post-depletion, Figure 4). This was done for both model (1) (Figure 4A) and model (2) (Figure 4B). The trends are qualitatively similar for both models. CTL depletion is predicted to result in an increase in the number of virus-producing cells, with the increase being more pronounced in the extra-follicular compared to the follicular compartment. More generally, the rise is more pronounced if the baseline number of virus-producing cells is lower due to a stronger CTL response. The increase in the number of virus-producing cells in the extra-follicular compartment is predicted to be stronger for model 2 (Figure 4A,B), because suppression of EF virus load is stronger for model (2) compared to model (1).

**Figure 4.**
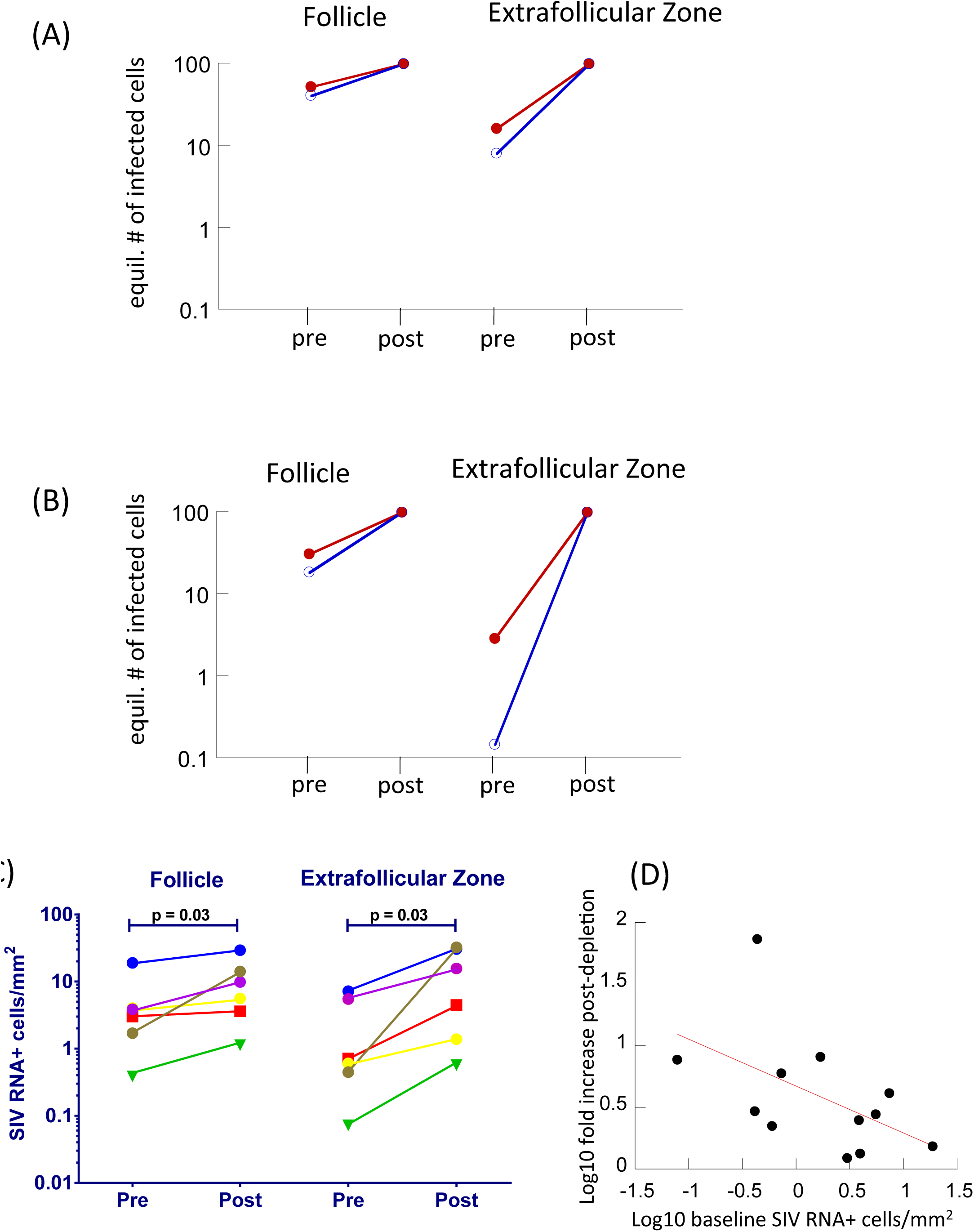
(A,B) Simulation of CTL depletion experiments with (A) model (1) assuming no antigen-induced CTL expansion in the follicular compartment, and (B) model (2) assuming the presence of antigen-induced CTL expansion in the follicular compartment. The equilibrium number of virus-producing cells is plotted in the presence of CTL (pre) and in their absence (post). This was done for a weaker CTL response / higher baseline virus load (red) and a stronger CTL response / lower baseline virus load (blue). Parameters were chosen as follows. λ_e_= λ_f_ =50, d=0.01, β_e_=β_f_=0.00072, a=0.45, η=0.01, b=0.01, p=0.001, g=0.5, h=2. For the weaker CTL response, c_e_=1 (and c_f_=1 for model 2), for the stronger CTL response c_e_=3 (and c_f_=3 for model 2). (C) Results from CD8 cell depletion experiments, performed in SIV-infected rhesus macaques, showing SIV RNA levels pre and 14 days post administration of an anti-CD8 antibody. The different colors represent data from different experimental animals. See text for details. The data are based on experiments generated for a previous study [27], but contain so far unpublished data. CD8 cell depletion results in a significant increase in the number of SIV RNA+ cells in both compartments, as shown. The fold-increase is significantly larger in the EF compared to the F compartment (p=0.03, Wilcoxon test). (D) Correlation between baseline SIV RNA levels pre depletion, and the fold-increase of SIV RNA levels post depletion. Data from both compartments are included, showing a significant negative correlation (p=0.04), as suggested by the mathematical models.

These predicted patterns are also seen in the experimental data. Although comparison to data cannot be used to distinguish between the two models, it does establish that the modeling approaches used here are consistent with available experimental information. We evaluated the impact of CD8+ cell depletion on the frequency of virus-producing cells within follicular and extrafollicular compartments in lymph nodes from 6 chronically SIV-infected rhesus macques as previously described [27]. Prior to CD8+ cell depletion, all animals demonstrated higher concentrations of virus-producing cells within the follicles than in the extrafollicular compartments. While in the current data, this difference does not reach statistical significance due to small sample size, a convincing statistical difference has been demonstrated in previous studies, both in rhesus macaques [12] and in humans [11,28]. Fourteen days after administration of an anti-CD8 antibody, frequenices of virus-producing cells increased in both compartments, and this increase was significantly more pronounced in the extrafollicular compartment, such that similar levels were found in both compartments. Notably, animals with the lowest frequencies of virus-producing cells at baseline demonstrated the largest increases in virus replication in both compartments, whereas those with the highest frequencies of virus-producing cells at baseline demonstrated more muted responses to CD8 depletion (Figure 4C, D). These findings further underscore the hypothesis that the CTL response is a major contributor to the distribution of virus within the lymph node.

#### CTL addition experiments

To address the deficiency of virus-specific CTL in B cell follicles and reduce virus replication at those sites, we have proposed to introduce the follicular homing molecule CXCR5 into virus-specific CTL [29]. Binding of CXCR5 by its ligand CXCL13 does not induce proliferation of T cells. It induces chemotaxis of the cells towards higher concentrations of CXCL13, which are in the follicle. It is possible that when the CTL contact virus-producing cells in the follicle, they will be stimulated to proliferate by engagement of the T cell receptor.

We simulated potential experiments where F-homing CTL are added to experimental animals. To do so, we considered a second population of CTL with F-homing characteristics and added those to the F compartment where cell populations were at equilibrium (details given in the legend to Figure 5). In this model, resident and added CTL were tracked as two distinct populations. The analysis was done for model (1) (Figure 5A), and for model (2) (Figure 5B). In terms of model parameters, the resident and added CTL populations differed as follows. The resident CTL were assumed to have a low migration rate from EF to F, but a high migration rate from F back to EF, as has been assumed so far. This results in the majority of the CTL residing in the extra-follicular compartment. The added CTL population has the opposite characteristics because they are supposed to be F-homing. They have a fast migration rate from EF to F, and a slow migration rate from F back to EF. Hence, the added CTL tend to accumulate in the F compartment. It was further assumed that the antigen-induced proliferation rate of the added CTL was somewhat lower than that of the resident CTL. The rationale behind this assumption is that experimental addition of CTL might result in a reduction in their efficacy compared to native CTL, although model results do not depend on this assumption.

**Figure 5.**
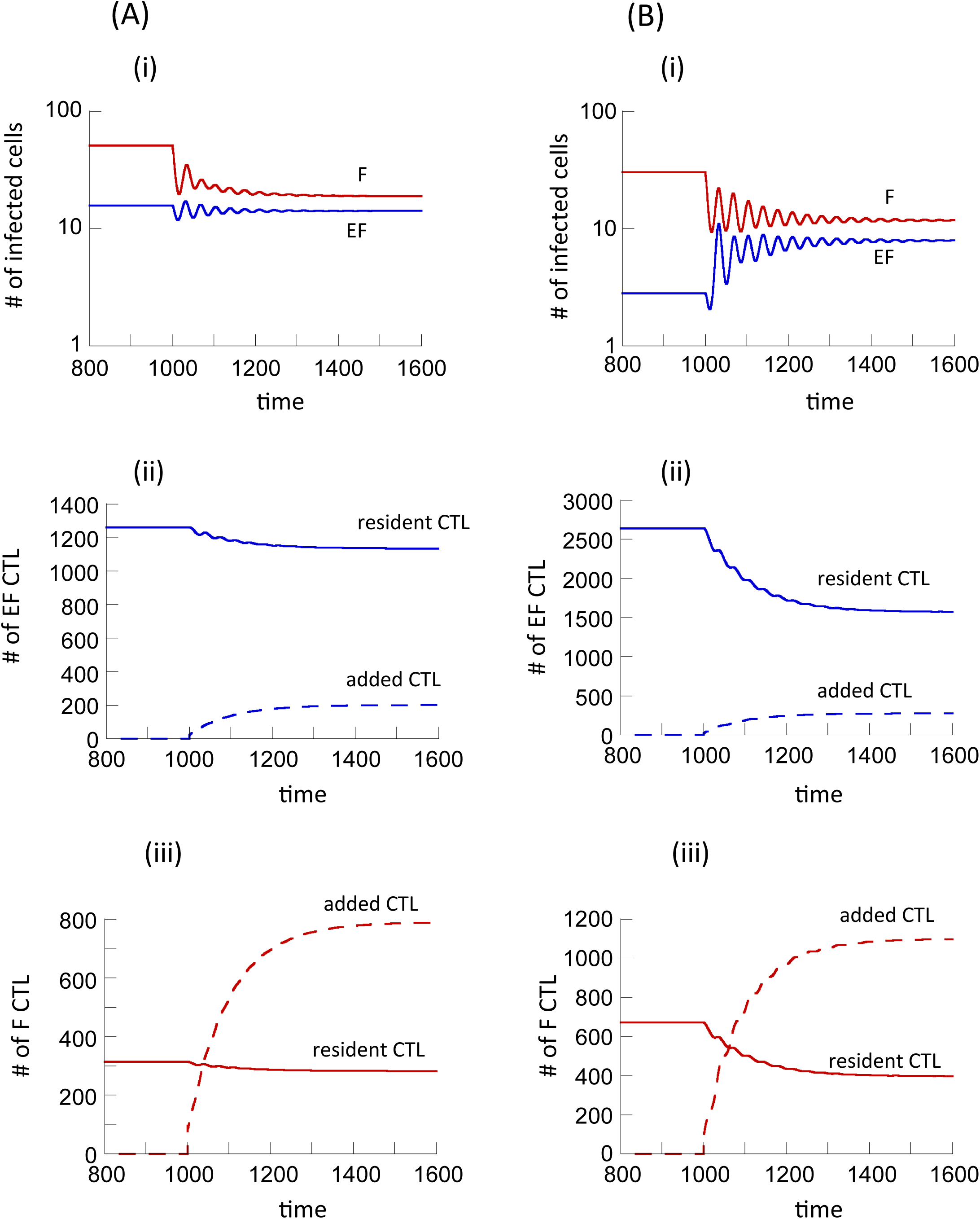
Simulation of experiments where CTL that preferentially home to the follicular compartment are added to a system at equilibrium, using (A) model (1) assuming no antigen-induced CTL expansion in the follicular compartment, and (B) model (2) assuming the presence of antigen-induced CTL expansion in the follicular compartment. The system was first allowed to equilibrate, and at time step 1000, one hundred F-homing CTL were added to the F compartment. Resident and F-homing CTL were tracked as separate populations, as described in the text. Both part A (left side) and part B (right side) have panels i-iii. Shown are the dynamics of virus-producing cells in the two compartments (panel i), the CTL dynamics in the EF compartment (panel ii), and the CTL dynamics in the F compartment (panel iii). Parameter values were chosen as follows. λ_e_= λ_f_ =50, d=0.01, β_e_=, β_f_=0.00072, a=0.45, η=0.01, c_e_=c_f_=1, b=0.01, p=0.001. For the resident CTL, g=0.5, h=2. For the added F-homing CTL, g=2, h=0.5. Further, we assumed a lower expansion rate for the added CTL, given by c2_e_=0.7 and also c2_f_=0.7 for model (2).

For Model 1, the simulation results are shown in Figure 5Ai. CTL addition results in a decline in the number of virus-producing cells in both compartments, with the decline being more pronounced in the follicular compartment. The virus-producing cell population declines only slightly in the extrafollicular compartment (which might be difficult to measure experimentally) because some of the added F-homing CTL enter the extrafollicular compartment through migration, resulting in additional killing there. This is also reflected in Figure 5Aii-iii, showing the dynamics of resident and added CTL in both compartments. The added F-homing CTL become dominant in the follicular compartment, and make up a minority population in the extrafollicular compartment.

The same analysis was performed for model (2) assuming that antigen-induced CTL expansion can occur in the follicular compartment. Now, slightly different behavior is observed (Figure 5Bi). As before, follicular virus load is reduced upon addition of the F-homing CTL. At the same time, however, extra-follicular virus load increases, which is the opposite compared to model (1). The reason is that in model (2), follicular virus stimulates CTL to proliferate, and the resident CTL resulting from this proliferation migrate back to the EF compartment with a fast rate, suppressing EF virus load. When F-homing CTL are added, this force is reduced. F-homing CTL reduce virus load in the follicular compartment and become dominant relative to the resident CTL in this site (Figure 5iii). Therefore, stimulation and subsequent migration of resident CTL back to the EF compartment occurs with a reduced rate, allowing the extrafollicular virus population to replicate at higher levels.

These predictions show that experiments involving the addition of F-homing CTL can help distinguish between the two models, based on the virus dynamics in the extra-follicular compartment that occurs upon addition of the F-homing CTL. While such experiments have not yet been reported, they are an important next step to perform. This knowledge will further be important to evaluate the use of F-homing CTL as a potential therapeutic strategy in order to reduce the relatively uncontrolled follicular virus population. If this indeed results in a rise in extrafolliuclar virus load, additional approaches would have to be employed to address this issue.

### Parameters and assumptions

The analysis concentrated on the effect of two key parameters: The rate of CTL expansion, c, and the rate at which CTL move from the extrafollicular to the follicular compartment, g.

In the literature, the rate of CTL expansion has often been used as a key parameter that determines the extent of CTL-mediated virus control. Other parameters, however, also drive virus load in our models. In the model formulations considered here, the rate of CTL-mediated killing, p, has the same effect as the rate of CTL expansion, c, such that the equilibrium virus load is determined by the product c*p. Therefore, the effect of the parameter p on the degree of virus compartmentalization is the same as that of the parameter c. Another important determinant of virus control is the death rate of CTL, b, with lower values of b resulting in lower equilibrium virus loads. Therefore, lower values of b result in more virus compartmentalization, as shown in Supplementary Text S1.

In the context of CTL migration processes, we varied the rate at which CTL move from the extrafollicular to the follicular compartment (parameter g) and kept the rate of back-migration from F to EF (parameter h) constant. The distribution of CTL can, however, can also be influenced by the parameter h. In fact, if the parameter g is sufficiently large, an increase in g has an identical effect as a decrease in h, and it is the ratio of g/h that counts (Supplementary Text S1). For lower value of g, however, the number of CTL that enter the F compartment is limited, and in this parameter regime, an increase in the parameter g is not quantitatively equivalent to a decrease in h and we need to consider the dependence of the outcome on the individual parameters rather than on the ratio of g/h. The reason is that if there are hardly any CTL in the follicular compartment (due to small value of g), variation in the migration rate from the follicular to the extra-follicular compartment does not have a large effect on the dynamics. This is illustrated with computer simulations in Supplementary Text S1.

In the simulations presented in the figures, parameter values were chosen such that the migration rates of infected cells were generally slower than those of CTL among the two compartments. As far as we are aware, there are currently no data available that would allow us to estimate the respective migration rates. The results presented in the figures, however, largely do not depend on those parameter choices (see Supplementary Text S1). If the migration rate of infected cells is sufficiently large, the degree of virus compartmentalization is reduced because the system essentially starts behaving more like a single well-mixed compartment, which is not a parameter region of interest in the current study.

Due to lack of appropriate data, it is further not clear how the number of target cells compares in the two compartments, and how cell parameters differ between the two compartments. For simulations, we chose parameters such that the number of susceptible target cells in the absence of infection is the same for the two compartments, but the results presented here do not depend on this choice of parameters. In Supplementary Text S1, we show that simulation results remain qualitatively the same if there are significant differences in target cell parameters and target cell numbers in the two compartments, brought about by differences in parameters λ_e_ and λ_f_. Results further remain robust if we assume asymmetrical migration rates of the target cells between the two compartments, and if CTL expansion occurs at different rates in the two compartments (Supplementary Text S1).

## Discussion and Conclusion

Experiments with SIV-infected macaques indicated that in chronic infection, there is an unequal distribution of virus in the follicular and extrafollicular compartment of lymphoid tissues [12,13]. The majority of the virus population was shown to reside in the follicular compartment. Data further indicate that CTL have a reduced ability to home to the follicular compartment, which can contribute to the observed virus compartmentalization. Since the differential CTL activity can contribute to the unequal virus distribution, the degree of virus compartmentalization is highest for strong CTL-mediated virus control (leading to low virus loads), and the distribution of the virus among the compartments becomes more equal for less efficient CTL-mediated virus control (higher virus loads) [12,13]. We used mathematical models to investigate in more detail how the dynamics of CTL responses can affect the distribution of the virus population across the two compartments, and how different assumptions about the CTL responses can influence outcome. In particular, there remains considerable uncertainty about CTL-related processes that occur in the follicular compartment. Data indicate that a relatively small population of CTL does enter the follicular compartment, and that these CTL can reduce virus load to a certain extent in this site [27]. It is unclear whether virus-producing cells in the follicular compartment are able to induce proliferation of the CTL, or whether expansion processes are absent in this compartment, and if so why. Therefore, we compared the properties of two models, one in which no CTL expansion was allowed to occur in the follicular compartment, and one in which expansion processes did take place there, with subsequent migration of the expanded CTL back to the extra-follicular compartment. The predictions of both models agreed with experimental data that are available so far. Thus, as seen in data, both models predict an increasing ratio of F:EF virus load with stronger CTL responses and lower equilibrium virus loads. In other words, while significant virus compartmentalization occurs in the model for strong CTL responses and low virus loads, virus distribution is predicted to be significantly more equal for weaker CTL responses and higher virus loads. Similarly, simulation of CTL depletion experiments with both models predicts that depletion results in a rise of virus load in both compartments, and that this rise is more pronounced in the extra-follicular compared to the follicular compartment, consistent with our experimental observations shown here and reported previously [27]. These two predictions are actually not independent of each other, as they both depend on the strength of the CTL response, and the models considered here might not be the only ones with these properties. The match of these predictions with experimental data, however, means that both model (1) and model (2) are consistent with available data, and hence are for now appropriate to explore.

While both models are consistent with available experimental observations, they significantly differ in other aspects. In particular, the degree of compartmentalization is predicted to be stronger for model (2) with follicular CTL proliferation compared to the model (1) without. In fact, in model (2), a possible outcome is the clearance of extra-follicular productive infection, were any virus-producing cells found in the extra-follicular compartment are the result of immigration from the follicle. There is experimental indication that virus-producing cells found in the EF compartment are indeed the result of immigration from the F compartment rather than the result of sustained extrafollicular virus replication, supporting the prediction of model (2) that productive infection in the EF compartment is essentially cleared by CTL: In the gut of SIV-infected macaques, it was observed that virus-producing cells in the extra-follicular region tended to be located near the follicles, in fact adjacent to them [12], indicating that they have recently immigrated from there. Additionally, this model-predicted outcome is supported by the data from well-controlled SIV infection, in which effective clearance of productive infection in the extra-follicular compartment was reported [12,30]. It is intriguing that Ki67+ virus-specific CTL are detected in follicles [27], adding to the data that support follicular CTL expansion as a physiologically relevant process, as assumed in model (2). In model (1) without follicular CTL expansion, on the other hand, there is no equilibrium or outcome that corresponds to clearance of extra-follicular productive infection. If the CTL response is very strong, the number of productively infected cells predicted by the model can in theory be sufficiently low to essentially correspond to clearance in a stochastic setting. Whether this is a likely scenario, however, is unclear, especially given data that document significant functional impairment of HIV/SIV specific CD4 T helper cell responses [31], which in turn negatively impact the effectiveness of CTL responses [5,32].

Our analysis indicates that experiments involving the addition of follicular-homing CTL to SIV-infected macaques could help to further distinguish between the two models considered here, based on the dynamics of virus-producing cells in the extra-follicular compartment following the addition of those CTL. If a significant increase of extra-follicular virus load occurs after CTL addition, CTL stimulation is likely to occur in the F compartment, as mathematically described in model (2). In contrast, if the addition of follicular-homing CTL does not result in significant changes in the number of extra-follicular virus-producing cells, we can discard the assumption that CTL can become stimulated in the follicle. If this is the case it is possible that other, yet unknown, mechanisms need to be invoked to explain the observed compartmentalization patterns and the effective clearance of productive infection in the extra-follicular compartment.

The use of chimeric antigen receptor T cells that express the follicular homing molecule CXCR5 could be an interesting therapeutic option to reduce virus load in the follicular compartment and hence to improve the overall degree of virus control. This could be a component of treatment approaches that aim for a cure. In this context, it will be important to further address the dynamics suggested by our models, in particular whether improved follicular virus control mediated by follicular homing CTL might lead to a concomitant increase in extrafollicular virus load as suggested by model (2).

As with any model, the results can depend on assumptions and the particular way in which the model is formulated. The biggest uncertainty of such models is the way in which the CTL response is formulated [17,33,34]. Many models used in the literature assume that the rate of CTL expansion is proportional not only to the number of virus-producing cells, but also to the number of CTL that are present. The dynamics in such models, however, tend to be rather unstable, involving pronounced and prolonged oscillations in CTL and virus populations, which are not typically observed in vivo. We chose a model where the rate of CTL expansion was only proportional to the number of virus-producing cells, because such models have been shown previously to display more realistic and stable behaviors that are more consistent with observations [17]. Nevertheless, computer simulations indicate that the main results reported here do not depend on these particular details.

Another aspect to note is that while the models considered here are consistent with currently available experimental data, this does not mean that the models are necessarily correct. There could be other models, with alternative assumptions that we have not thought of, that are just as consistent with the available experimental data. On the flip side, if a mathematical model is not consistent with data, we can discard the underlying assumptions with some confidence. For example, one mechanism that we hypothesized might independently drive infected cell compartmentalization is a difference in the activation status of CD4 T cells in the two compartments. While follicular CD4 T cells are by default activated and highly permissive to infection [35], the activation status and thus permissiveness of extra-follicular CD4 T cells might vary, and might be driven by the amount of virus present in the EF compartment [36]. We formulated such a process mathematically in Supplementary Text S1, and concluded that this mechanism alone does not result in dynamics that are consistent with data available so far.

While there are uncertainties about particular aspects of the models, they clearly demonstrate how differences in assumptions about the mechanisms of CTL activity in the two compartments can lead to different dynamics. This in turn can help to interpret future experimental data, to potentially exclude certain hypotheses about CTL activity in the follicular compartment, and to sharpen our thinking about how potential therapeutic efforts to increase CTL-mediated activity in the follicular compartment could impact the overall dynamics of the infection.

## Methods

The majority of this paper is computational in nature. The mathematical models and the analysis are fully presented in Results section as well as in Supplementary Text S1. In addition to the computational analysis, experimental data from CD8 cell depletion experiments in SIV-infected macaques are presented. These data were generated in the context of a previous study, according to the same methodology described in [27]. Procedures are briefly outlined in the manuscript text. The data shown in figures have, however, not been previously published.

## Supplementary Text S1

### Parameter dependencies and scaling

The analysis presented in the main text concentrated mainly on varying two parameters: The parameter c, the rate of CTL expansion, was varied because it is a major determinant of the equilibrium number of infected cells, and hence of the degree of CTL-mediated virus control. It is therefore a measure of the CTL “strength”. The parameter g was varied because we wanted to explore the effect of changing the ability of the CTL to home to the follicular compartment. Besides the parameters that we varied in the main part of our analysis, however, other parameters also influence the ability of CTL to reduce virus load, or the tendency of the CTL to accumulate in the follicular compartment. In section 5 of the main text, we discuss the effects of those alternative parameters. In the current section, we present select simulation results to support the arguments in the main text. Further, due to lack of estimates for some of the parameter values, arbitrary assumptions were made. Thus, we assumed that the number of susceptible target cells was identical in the two compartments, and that the movement rate of infected cells from the follicular to the extrafollicular compartment is the same as that for the opposite direction. In section 5 of the main text, we re-visited predictions in parameter regimes where target cell kinetics differ in the two compartments, and where movement rates between the two compartments are asymmetric. Supporting computer simulations are shown in the current section.

#### Varying the degree of CTL-mediated virus control

As mentioned in the main text, besides the rate of CTL expansion, c, the equilibrium number of infected cells is also determined significantly by other CTL parameters, such as the rate of CTL-mediated lysis, p, and the death rate of CTL. b. In fact, the parameter p has an identical effect on the equilibrium number of infected cells as the parameter c. That is, a given increase the parameter c results in a reduction in the equilibrium number of infected cells that is identical to that observed for the same increase in the parameter p (not shown). Therefore, these two parameters are interchangeable in the context of the model we investigated here.

Due to unknown immune parameter values, the figures shown in the main text for model (2) assume identical CTL expansion rates for the extrafollicular and follicular compartments, i.e. c_e_=c_f_. Here, we show that results remain similar if we assume different CTL expansion rates in the two compartments. In Figure S1, the graph from Figure 3A in the main text is replotted in black (c_f_ = c_e_), in addition to an equivalent graph assuming that c_f_ = 0.5c_e_ (Red). This demonstrates that the general patterns remain qualitatively the same.

**Figure S1:**
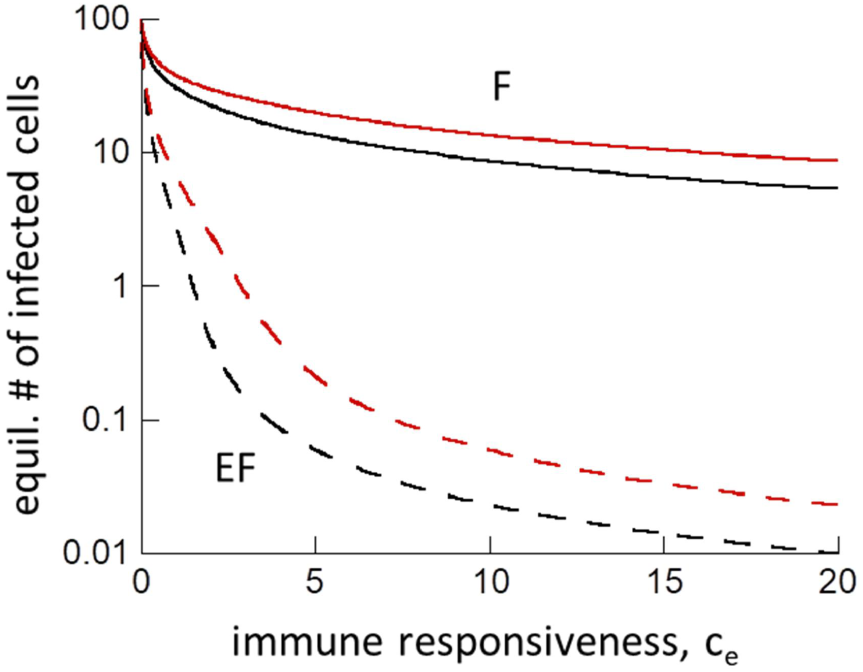
(A) Equilibrium number of virus-producing cells in the two compartments as a function of the immune responsiveness c_e_. The black lines show the same graph as in Figure 3A in the main text (c_f_=c_e_), and the red lines are new, assuming c_f_=0.5c_e_. F=follicular compartment (solid lines), EF=extrafollicular compartment (dashed lines). Parameters are the same as in Figure 3A in the main text.

The CTL death rate, b, is also an important determinant of the number of infected cells at equilibrium, with lower values of b (longer lived CTL) resulting in fewer infected cells. Figure S2A plots the equilibrium number of infected cells in the two compartments as a function of 1/b, and the dependence is qualitatively the same compared to the dependence on the CTL expansion rate, c (compare to Figure 2A in main text). Figure S2B uses these data to plot the total number of infected cells against the F:EF ratio, and the pattern is again qualitatively the same compared to varying the parameter c (Figure 2B in main text).

**Figure S2:**
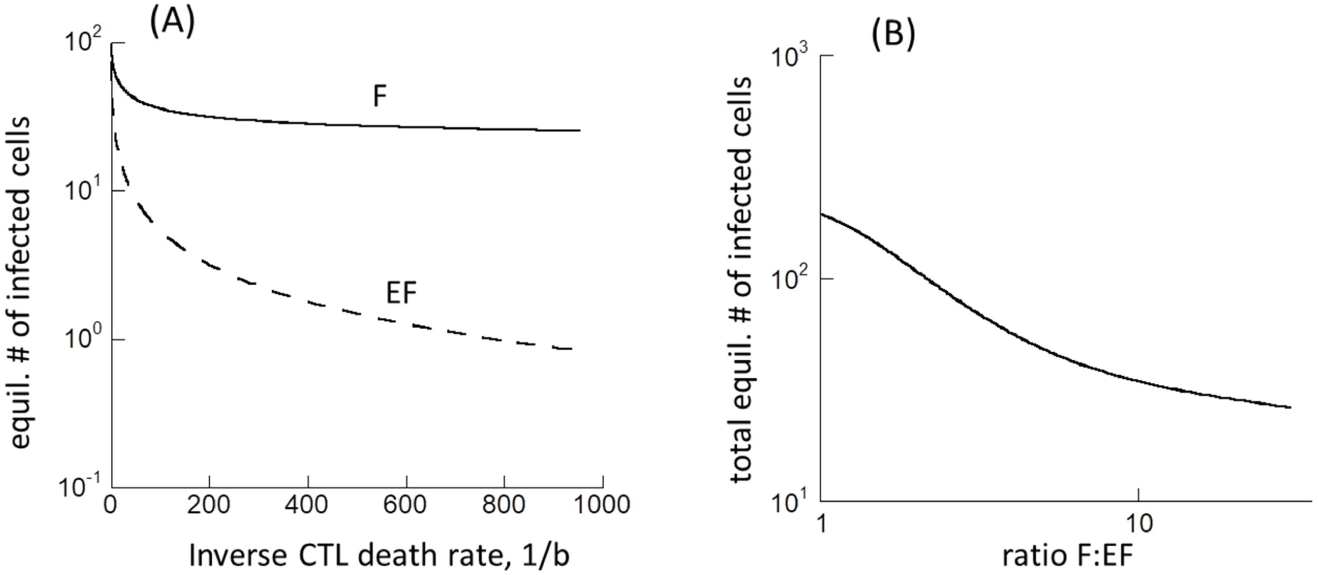
(A) Equilibrium number of virus-producing cells in the two compartments as a function of 1/b. F=follicular compartment, EF=extrafollicular compartment. (B) Relationship between the equilibrium number of virus-producing cells and the ratio of follicular to extra-follicular virus load (F:EF). Parameters were the same as in Figure 2A, but with c=5.

#### Varying the rate of CTL homing to the follicular compartment

In the main part of our analysis, we varied the rate of CTL homing from the extrafollicular to the follicular compartment, g. An increase in the parameter g corresponds to increased homing to the follicular compartment, and hence to more CTL being present in that site. This process is countered by the rate of CTL homing away from the follicular compartment, towards the extrafollicular site, h. If the parameter g is sufficiently large, a decrease in the parameter h has an identical effect compared to an increase in g. Therefore, it is really the ratio of g/h that determines outcome. This is shown in Figure S3, comparing the black and red lines. The black line is the same graph shown in in the main text in Figure 2C, where only the parameter g was varied. The red line represents the outcome of corresponding simulations where the parameter h was varied. For lower values of g, however, we find that a decrease in h leads to different results compared to an increase in g. Hence, it is not the ratio g/h that determines outcome (Figure S3, compare blue and black lines). The reason is that for relatively low values of g, comparatively few CTL enter the follicular compartment. A reduction in the rate of CTL outflux from the follicular compartment, h, therefore is not identical to an increase in the rate of CTL influx, g. If there are hardly any CTL in the follicular compartment, a reduced rate at which they leave this site cannot increase the number of follicular CTL by a large amount.

**Figure S3:**
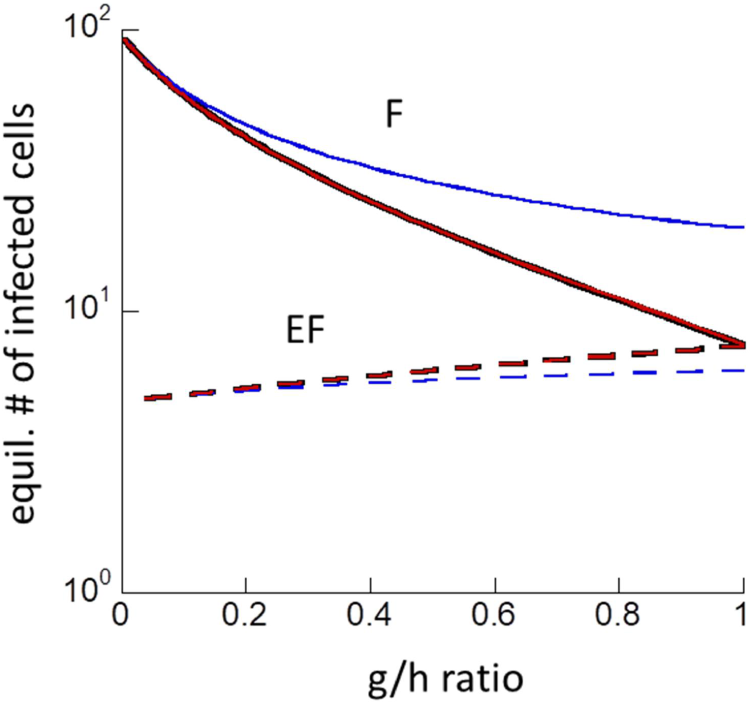
Equilibrium number of virus-producing cells in the two compartments as a function of the ratio g/h. The black lines are identical to the ones in Figure 2C, and were generated by varying the parameter g. The red lines were generated by varying the parameter h, assuming a relatively large value of g (g=1). The blue lines were also generated by varying the parameter h, but assuming a lower value of g (g=0.01). The remaining parameter values were the same as in Figure 2C.

#### Effect of infected cell migration rate

In the computer simulations presented in the main text, a relatively low rate of migration of infected cells between the compartments, η, was assumed. We also explored the outcome when the migration rate of infected cells was increased, to an order of magnitude that is equivalent to the CTL migrations rates that were assumed in those simulations. Results remained qualitatively the same, although the extent of compartmentalization was reduced for larger values of η (Figure S4).

**Figure S4:**
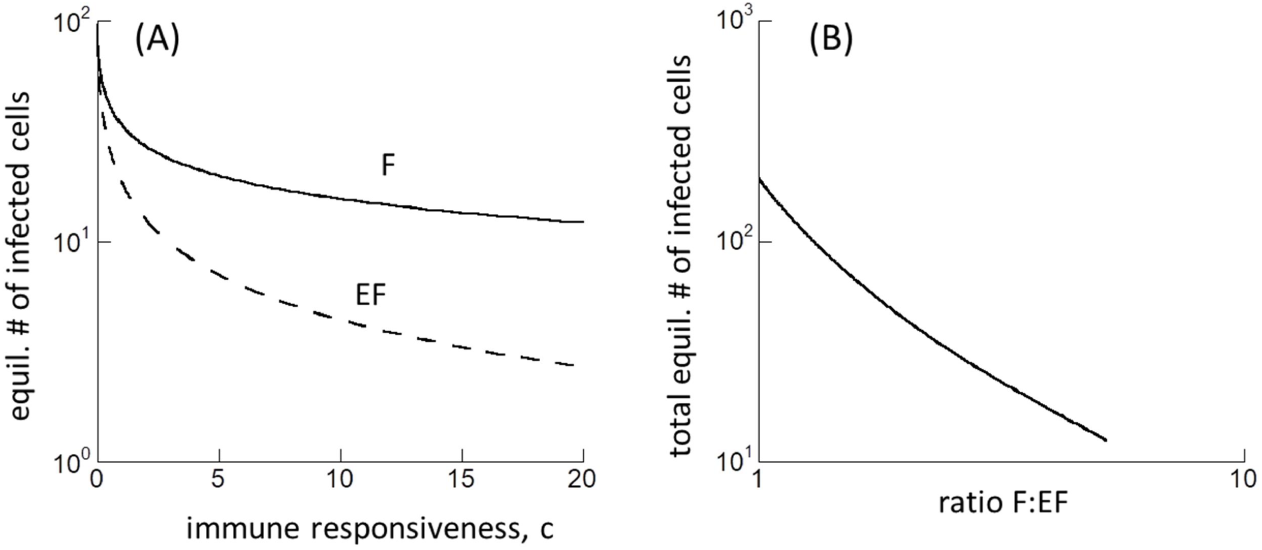
Same simulations as in Figure 2A and 2B in the main text, but with a larger infected cell migration rate η=0.5. (A) Equilibrium number of virus-producing cells in the two compartments as a function of the CTL responsiveness, c, which correlates with the strength of the CTL response. F=follicular compartment, EF=extrafollicular compartment. (B) Relationship between the equilibrium number of virus-producing cells and the ratio of follicular to extra-follicular virus load (F:EF).

If the rate of infected cell migration, η, is increased to sufficiently high levels, the degree of compartmentalization becomes very small (Figure S5). Effectively, the system behaves more like a well-mixed pool of cells.

**Figure S5:**
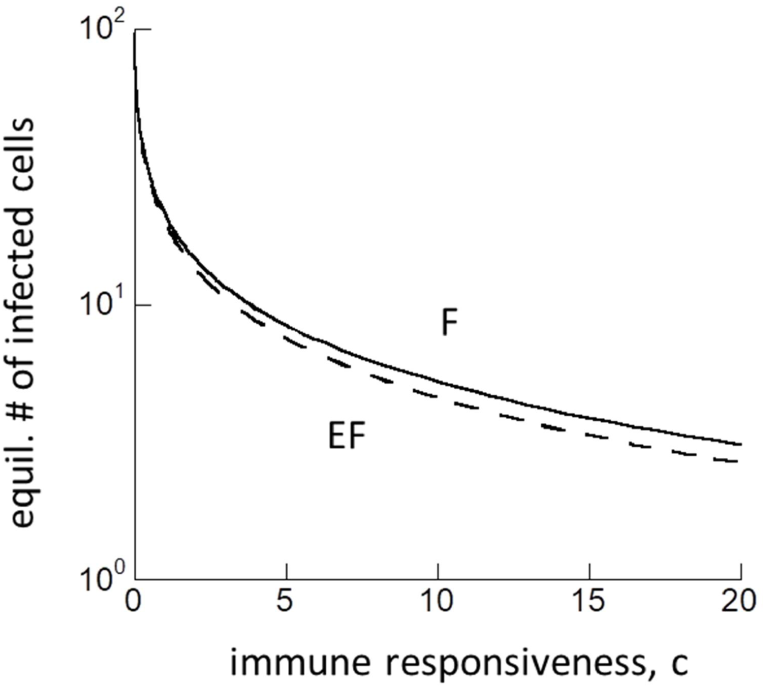
Same simulations as in Figure 2A, but with a large infected cell migration rate η=10. The equilibrium number of virus-producing cells in the two compartments is plotted as a function of the CTL responsiveness, c, which correlates with the strength of the CTL response. F=follicular compartment, EF=extrafollicular compartment.

#### Compartment-specific parameters

So far, all simulations assumed that the target cell kinetics were the same in the two compartments, i.e. that the number of target cells in the absence of infection was the same in the follicular and extrafollicular compartments. Further, it was assumed that infected cells migrated with equal rates in both directions. Here, were repeat the simulations in Figure 2A and B from the main text assuming that target cell numbers are different in the two compartments in the absence of infection, and that migration rates are different in each direction. Thus, we assume that the values of λ_e_ and λ_f_ were different. The migration rate of infected cells from the extrafollicular to the follicular compartment is now denoted by η_1_, while the migration rate in the opposite direction is denoted by η_2_. As shown in Figure S6, the general dependencies remain the same compared to the ones presented in Figure 2A/B in the main text.

**Figure S6:**
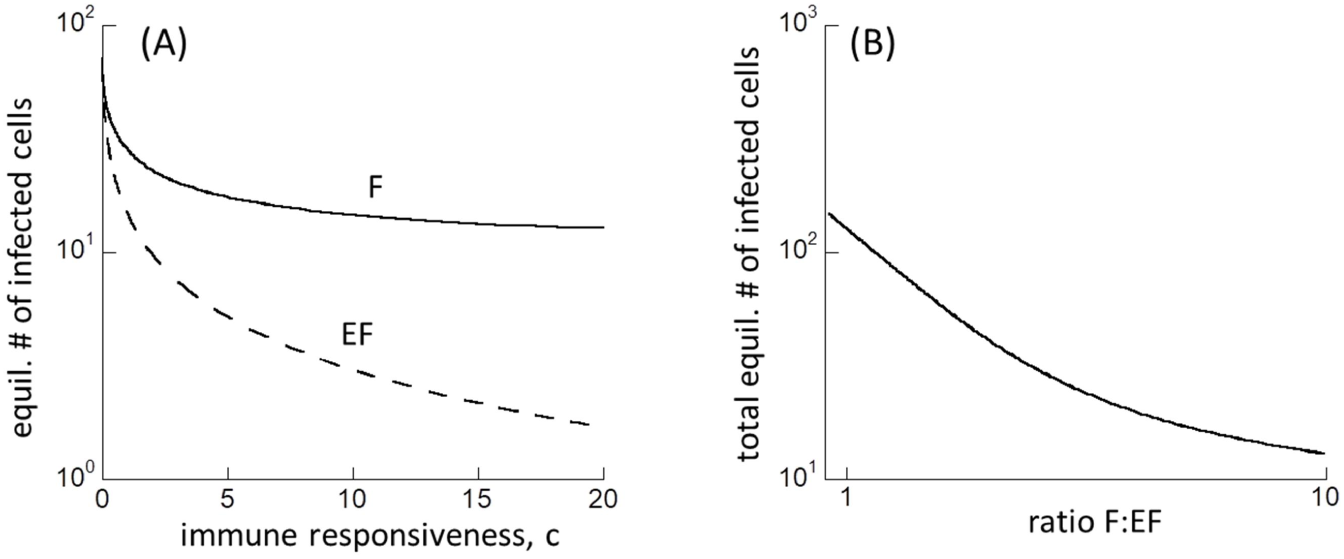
Same simulations as in Figure 2A and 2B in the main text, but with λe=50 and λf=30, as well as η1=0.1 and η2=0.01. F=follicular compartment, EF=extrafollicular compartment.

**Target cell activation, infection permissiveness, and compartmentalization**

While CD4 T cells in the follicular compartment are always activated and by definition highly permissive to infection, in the extra-follicular compartment, target cell permissiveness might depend on antigen-induced stimulation of the T cells [1]. In other words, the permissiveness of the T cells to infection might depend on virus load in the extra-follicular compartment. This could in principle be another mechanism that drives the observed compartmentalization of infected cells, and we tested this hypothesis with a mathematical model. To do so, model (1) in the main text was modified to assume that productive target cell infection requires their activation in the extra-follicular compartment, and that activation occurs in response to antigenic stimulation by the virus. For the dynamics in the follicular compartment, we assumed that all target cells are activated and permissive to infection. Denoting resting (non-susceptible) target cells in the extra-follicular compartment by W_e_, and activated target cells as X_e_, the model is thus given by the following set of ordinary differential equations.

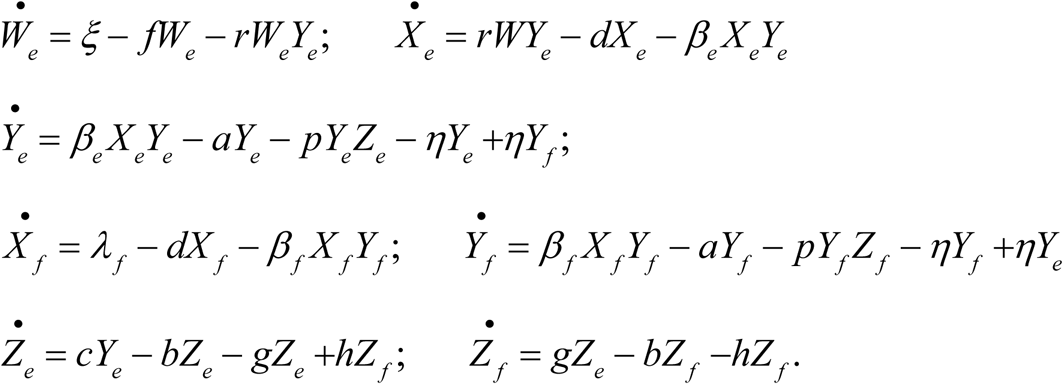

Resting target cells are produced with a rate ξ, die with a rate f, and become activated by the presence of productively infected cells with a rate r.

In the context of a single compartment, virus dynamics under the assumption that target cell permissiveness depends on virus-induced cell activation has been studied before [2], and the properties are briefly outlined here. This kind of model is characterized by two equilibria being simultaneously stable: the virus extinction equilibrium, and a virus persistence outcome. To which equilibrium the system converges is determined by initial conditions, in particular initial virus load. If initial virus load is relatively high, enough stimulation of target cells occurs to generate a sufficient number of activated cells to enable persistent infection. If initial virus load is low, however, not enough stimulation is present, which fails to generate sufficient numbers of activated target cells for sustained replication. As a consequence, the population of virus-producing cells goes extinct. This extinction is further promoted by the presence CTL responses, which can suppress virus load to a level that is too low to maintain a sufficient level of cell activation. The same kind of dynamics can occur in our current model and we focus parameters in which the virus population is essentially driven extinct in a one-compartment setting due to lack of target cell activation. The model discussed here, however, takes into account a second compartment (follicular compartment), from which infected cells can enter by migration. The infected cells that migrate to the extra-follicular compartment can also contribute to target cell stimulation. Hence, the outcome depends on the rate of infected cell migration from the F to the EF compartment (parameter η), which is described as follows.

If the migration rate, η, is relatively high, the incoming infected cells activate a sufficient number of target cells in the EF compartment such that sustained infection in this compartment is possible (Figure S7 A). If the migration rate, η, is low, however, this is not the case anymore. In this scenario, no active virus replication occurs in the EF compartment, and only a small population of extra-follicular virus-producing cells remains, as a result of migration from the follicular compartment (Figure S7 C). For intermediate migration rates, η, dynamics are observed that lie between the two cases described above (Figure S7 B). The number of extra-follicular virus-producing cells declines towards extinction, since CTL push virus loads to levels that are too low to maintain enough activated target cells. Limited migration of virus-producing cells, however, periodically activates target cells, which allows the virus population to grow in the EF compartment and to further activate T cells, resulting in positive feedback-like dynamics and virus expansion (Figure S7 B). The CTL response, however, reacts to this and eventually pushes the virus population again to levels that are too low to sustain target cell activation. These dynamics are repeated periodically, resulting in stable population cycles over time (Figure S7 B).

**Figure S7:**
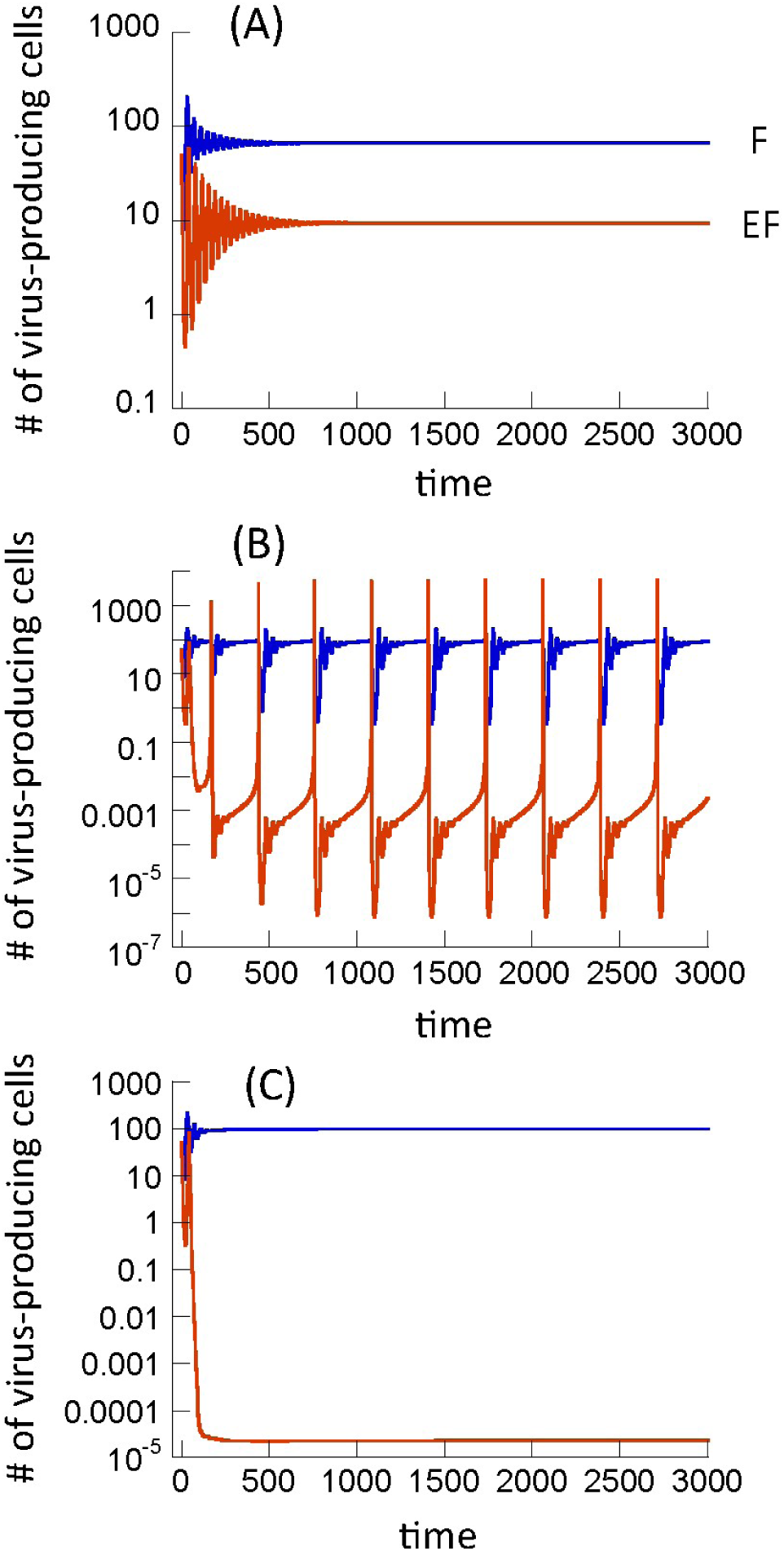
Dynamics in F and EF compartments observed in the target cell activation model. Parameters were chosen as follows. ξ= λ_f_ =50, f=0.001, d=0.01, β_e_=, β_f_=0.00072, a=0.45, c_e_=2, b=0.01, p=0.001, g=0.5, h=2. The panels differ in the migration rate of virus producing cells, η. (A) η=10^−3^, (B) η=10^−5^, (C) η=10^−7^.

The most relevant regime for our investigation is the one shown in Figure S7 C, where virus-producing cells in the extra-follicular compartment are not sustained by virus replication in that site, but by influx of cells from the follicular compartment. This results in stronger compartmentalization compared to model (1) in the main text, as shown in Figure S8. In contrast to the properties of model (1), however, the target cell activation model fails to predict a rise of virus load in the extra-follicular compartment following CTL depletion (Figure S8). The reason is that virus load in the EF compartment is too low to maintain a sufficient number of activated target cells in the presence of CTL. Once the CTL have been removed, there are still not enough activated target cells present to enable virus expansion, because a threshold amount of virus in the EF compartment is required to achieve this. Hence, without a sufficient external influx of target cell stimulation, an immediate growth of the virus population in the EF compartment following CTL depletion is not expected in this model. Therefore, the properties of this scenario are in conflict with the experimental data discussed in the main text.

**Figure S8:**
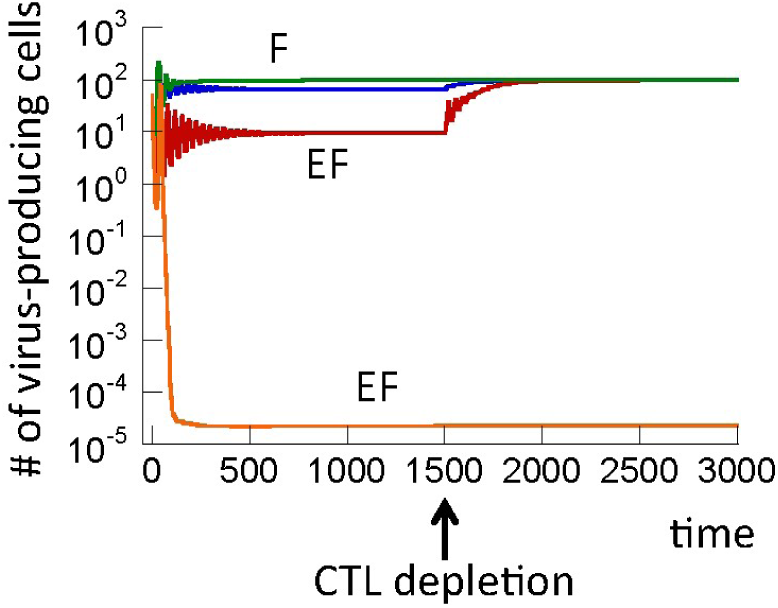
Comparing the pre- and post CTL depletion dynamics in model (1) from the main text (F=blue, EF=red) and in the target cell activation model in the regime depicted in Figure S1 C (F=green, EF=orange). For the target cell activation model, CTL depletion does not result in an increased virus load. See text for explanation. Parameters were chosen as follows. ξ= λ_e_ =λ_f_ =50, f=0.001, d=0.01, β_e_=, β_f_=0.00072, a=0.45, c_e_=2, b=0.01, p=0.001, g=0.5, h=2. For model (1), η=0.01. For the target cell activation model, η=10^−7^.

We now focus on the parameter regime shown in Figure S7A, where infected cell influx from the F compartment is sufficient to maintain enough activated target cells to allow persistent virus replication in the EF compartment. We asked whether in this scenario, the requirement of target cell activation for infection changes the compartmentalization dynamics in a significant way. Hence, we compare computer simulations of model (1) with those of the activation model, assuming that the rate of target cell production is identical in the two models (λ_e_=ξ). As shown in Figure S9, the equilibrium numbers of virus-producing cells in the two compartments are very similar to each other. The reason is that in these types of mathematical models, equilibrium virus load is mostly determined by parameters of the immune response, rather than by parameters that govern the target cell dynamics [3]. Hence, the models suggest that in this parameter region, the requirement of target cell activation for infection does not contribute significantly to the observed compartmentalization of the virus population.

**Figure S9:**
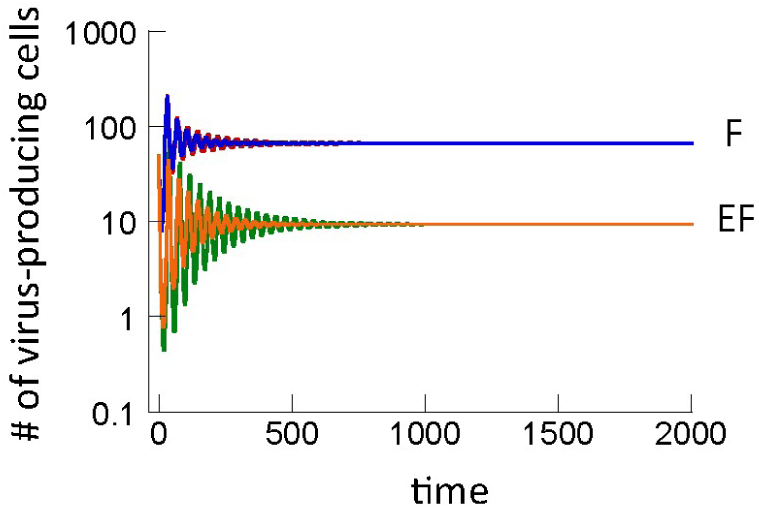
Comparison of the dynamics observed in model (1) (red and green) and in the target cell activation model for the regime depicted in Figure S1 A, where enough cell activation occurs to allow sustained infection in the EF compartment (blue and orange). In this case, there is no significant difference between the two models. Parameters were chosen as follows. *ξ = λ_e_ =λ_f_ =50, f=0.001, d=0.01, β_e_=, β_f_=0.00072, a=0.45, c_e_=2, b=0.01, p=0.001, g=0.5, h=2. For model (1), η=0.01. For the target cell activation model, η=10^−3^*.

Finally, we consider the parameter regime in Figure S7 B, where persistent population cycles are observed. We are not aware of data that document such cycles in the EF compartment, and hence the biological relevance of this regime is unclear. The occurrence of sustained population cycles is not the consequence of the two-compartment nature of the model. The same kind of dynamics are observed if we consider a single-compartment model with a constant external influx of virus-producing cells (not shown). It also continues to be observed if the model is altered in a number of ways, for example by introducing saturation terms into the rate of target cell activation or the rate of infection (not shown). The potential meaning of this parameter regime remains to be explored.

## References

1. Schmitz JE, Kuroda MJ, Santra S, Sasseville VG, Simon MA, et al. (1999) Control of viremia in simian immunodeficiency virus infection by CD8(+) lymphocytes [In Process Citation]. Science 283: 857–860.

2. Lifson JD, Nowak MA, Goldstein S, Rossio JL, Kinter A, et al. (1997) The extent of early viral replication is a critical determinant of the natural history of simian immunodeficiency virus infection. J Virol 71: 9508–9514.

3. Trachtenberg E, Korber B, Sollars C, Kepler TB, Hraber PT, et al. (2003) Advantage of rare HLA supertype in HIV disease progression. Nat Med 9: 928–935.

4. Chouquet C, Autran B, Gomard E, Bouley JM, Calvez V, et al. (2002) Correlation between breadth of memory HIV-specific cytotoxic T cells, viral load and disease progression in HIV infection. AIDS 16: 2399–2407.

5. Lifson JD, Rossio JL, Piatak M, Jr., Parks T, Li L, et al. (2001) Role of CD8(+) lymphocytes in control of simian immunodeficiency virus infection and resistance to rechallenge after transient early antiretroviral treatment. J Virol 75: 10187–10199.

6. Janeway C, P. T, M. W, D. CJ (1999) Immunobiology: The immune system in health and disease. New York: Current Biology Ltd.

7. Walker BD, Yu XG (2013) Unravelling the mechanisms of durable control of HIV-1. Nat Rev Immunol 13: 487–498.

8. Tenner-Racz K, Racz P, Bofill M, Schulz-Meyer A, Dietrich M, et al. (1986) HTLV-III/LAV viral antigens in lymph nodes of homosexual men with persistent generalized lymphadenopathy and AIDS. Am J Pathol 123: 9–15.

9. Embretson J, Zupancic M, Ribas JL, Burke A, Racz P, et al. (1993) Massive covert infection of helper T lymphocytes and macrophages by HIV during the incubation period of AIDS. Nature 362: 359–362.

10. Pantaleo G, Graziosi C, Demarest JF, Butini L, Montroni M, et al. (1993) HIV infection is active and progressive in lymphoid tissue during the clinically latent stage of disease. Nature 362: 355–358.

11. Connick E, Mattila T, Folkvord JM, Schlichtemeier R, Meditz AL, et al. (2007) CTL fail to accumulate at sites of HIV-1 replication in lymphoid tissue. J Immunol 178: 6975–6983.

12. Connick E, Folkvord JM, Lind KT, Rakasz EG, Miles B, et al. (2014) Compartmentalization of simian immunodeficiency virus replication within secondary lymphoid tissues of rhesus macaques is linked to disease stage and inversely related to localization of virus-specific CTL. J Immunol 193: 5613–5625.

13. Miles B, Connick E (2016) TFH in HIV Latency and as Sources of Replication-Competent Virus. Trends Microbiol 24: 338–344.

14. Nowak MA, May RM (2000) Virus dynamics. Mathematical principles of immunology and virology.: Oxford University Press.

15. Perelson AS (2002) Modelling viral and immune system dynamics. Nature Rev Immunol 2: 28–36.

16. Perelson AS, Ribeiro RM (2013) Modeling the within-host dynamics of HIV infection. BMC Biol 11: 96.

17. Wodarz D (2006) Killer cell dynamics: mathematical and computational approaches to immunology. New York: Springer.

18. Funk GA, Jansen VA, Bonhoeffer S, Killingback T (2005) Spatial models of virus-immune dynamics. J Theor Biol 233: 221–236.

19. Frost SD, Dumaurier MJ, Wain-Hobson S, Brown AJ (2001) Genetic drift and within-host metapopulation dynamics of HIV-1 infection. Proc Natl Acad Sci U S A 98: 6975–6980.

20. Perelson AS, Neumann AU, Markowitz M, Leonard JM, Ho DD (1996) Hiv-1 Dynamics in-Vivo -Virion Clearance Rate, Infected Cell Life-Span, and Viral Generation Time. Science 271: 1582–1586.

21. Ribeiro RM, Qin L, Chavez LL, Li D, Self SG, et al. (2010) Estimation of the initial viral growth rate and basic reproductive number during acute HIV-1 infection. J Virol 84: 6096–6102.

22. Nowak MA, Lloyd AL, Vasquez GM, Wiltrout TA, Wahl LM, et al. (1997) Viral dynamics of primary viremia and antiretroviral therapy in simian immunodeficiency virus infection. J Virol 71: 7518–7525.

23. Little SJ, McLean AR, Spina CA, Richman DD, Havlir DV (1999) Viral dynamics of acute HIV-1 infection. J Exp Med 190: 841–850.

24. Nowak MA, Bangham CR (1996) Population dynamics of immune responses to persistent viruses. Science 272: 74–79.

25. Holt RD (1977) Predation, Apparent Competition and the structure of prey communities. Theor Pop Biol 12: 197–229.

26. Bonsall MB, Hassell MP (1998) Population dynamics of apparent competition in a host-parasitoid assemblage. J Anim Ecol 67: 918–929.

27. Li S, Folkvord JM, Rakasz EG, Abdelaal HM, Wagstaff RK, et al. (2016) Simian Immunodeficiency Virus-Producing Cells in Follicles Are Partially Suppressed by CD8+ Cells In Vivo. J Virol 90: 11168–11180.

28. Folkvord JM, Armon C, Connick E (2005) Lymphoid follicles are sites of heightened human immunodeficiency virus type 1 (HIV-1) replication and reduced antiretroviral effector mechanisms. AIDS Res Hum Retroviruses 21: 363–370.

29. Skinner PJ, Connick E (2014) Skinner, P. J., and E. Connick. “Overcoming the immune privilege of B cell follicles to cure HIV-1 infection. J Hum Virol Retrovirol 1: 00001–00003.

30. Fukazawa Y, Lum R, Okoye AA, Park H, Matsuda K, et al. (2015) B cell follicle sanctuary permits persistent productive simian immunodeficiency virus infection in elite controllers. Nat Med 21: 132–139.

31. Douek DC, Brenchley JM, Betts MR, Ambrozak DR, Hill BJ, et al. (2002) HIV preferentially infects HIV-specific CD4+ T cells. Nature 417: 95–98.

32. Thomsen AR, Nansen A, Christensen JP, Andreasen SO, Marker O (1998) CD40 ligand is pivotal to efficient control of virus replication in mice infected with lymphocytic choriomeningitis virus. J Immunol 161: 4583–4590.

33. Wodarz D (2007) Killer Cell Dynamics: mathematical and computational approaches to immunology. New York: Springer.

34. Antia R, Ganusov VV, Ahmed R (2005) The role of models in understanding CD8+ T-cell memory. Nat Rev Immunol 5: 101–111.

35. Kohler SL, Pham MN, Folkvord JM, Arends T, Miller SM, et al. (2016) Germinal Center T Follicular Helper Cells Are Highly Permissive to HIV-1 and Alter Their Phenotype during Virus Replication. J Immunol 196: 2711–2722.

36. Cartwright EK, Spicer L, Smith SA, Lee D, Fast R, et al. (2016) CD8(+) Lymphocytes Are Required for Maintaining Viral Suppression in SIV-Infected Macaques Treated with Short-Term Antiretroviral Therapy. Immunity 45: 656–668.

## References

1. Cartwright EK, Spicer L, Smith SA, Lee D, Fast R, et al. (2016) CD8(+) Lymphocytes Are Required for Maintaining Viral Suppression in SIV-Infected Macaques Treated with Short-Term Antiretroviral Therapy. Immunity 45: 656–668.

2. Wodarz D, Lloyd AL, Jansen VA, Nowak MA (1999) Dynamics of macrophage and T cell infection by HIV. J Theor Biol 196: 101–113.

3. Wodarz D (2007) Killer Cell Dynamics: mathematical and computational approaches to immunology. New York: Springer.

